# Comprehensive landscape of potential CTCF-binding sites across the complete telomere-to-telomere human genome

**DOI:** 10.64898/2026.07.18.739380

**Authors:** Eleftherios Bochalis, Ilias Georgakopoulos-Soares

## Abstract

CCCTC-binding factor (CTCF) is a master regulator of the human genome, directly modulating both transcription and 3D genome organization. However, previous human genome-wide maps of its binding sites have been built on an incomplete assembly, leaving out many repetitive, difficult-to-sequence regions. Here, we leverage the telomere-to-telomere human assembly to generate a complete catalog of potential CTCF-binding sites and characterize their function in centromeric satellites, rDNA arrays, and acrocentric short arms. Using GC-stratified permutation testing, we find that CTCF motifs are significantly enriched in the alpha-satellite active higher-order repeats of chromosomes 18, 20, and X, in rDNA arrays, and in gamma satellites. Within acrocentric short arms, CTCF motifs concentrate in the hypomethylated and accessible regions of rDNA arrays and composite beta satellites. Although short arms share high sequence homology, most potential CTCF sites are arm-specific. Cross-arm shared sites were 1.7-fold more likely to fall in the frequently recombining pseudo-homologous regions, with density scaling with sequence mosaicism. Extending our analysis to non-human primate assemblies, we show that CTCF enrichment in gamma satellites and rDNA arrays is consistent across species. Subterminal heterochromatin spacer regions contain CTCF sites deriving largely from ancient LINE elements and lineage-specific segmental duplications act as a site transfer mechanism between species. We believe that our findings will provide the foundation for future studies of CTCF’s function in the repetitive parts of the human genome.

## Introduction

CCCTC-binding factor (CTCF) is a highly conserved zinc finger DNA-binding protein that functions as a key architectural regulator of the genome ^1^. It binds specific DNA motifs and, often in cooperation with cohesin, helps organize chromatin into loops and topologically associating domains. Pairs of convergent CTCF motifs anchor cohesin extrusion, defining the boundaries of these loops ^2^. Beyond its role in three-dimensional chromatin organization, CTCF contributes to promoter-enhancer interactions, transcriptional activation or repression, and the maintenance of epigenetic state at imprinted loci, such as the imprinting control region (ICR) of *Igf2* and *H19*, where CTCF binding regulates allele-specific expression ^1,3^.

CTCF contains a central zinc-finger domain, responsible for DNA recognition, flanked by N-terminal and C-terminal domains ^4^. The zinc-finger domain consists of 11 zinc-finger motifs (ZF) organized in a tandem array. ZF3-ZF7 bind to a core DNA sequence, whereas ZF8-ZF11 and ZF1-ZF2 clusters detect upstream and downstream non-conserved DNA sequences. When the flanking ZF clusters are utilized, an increase in binding affinity and chromatin residence time is observed ^5,6^. Since transcription factor binding is highly sequence-dependent, the sequence preferences of CTCF are commonly summarized in position weight matrices (PWMs) ^7^. This applies to other known transcription factors.

The epigenetic state of a DNA sequence directly affects the binding potential of CTCF. Cytosine methylation at CpG dinucleotides disrupts its binding ^8^. The core motif contains potential cytosine methylation sites at positions 2, 4, and 12, with methylation at position 2 reducing CTCF binding affinity by approximately 23-fold ^6,8^. A concrete example of how this methylation sensitivity has a direct functional consequence is the previously mentioned ICR of *Igf2* and *H19*. In this region, the ICR is methylated only on the paternal allele, enabling CTCF to bind to the corresponding unmethylated region in the maternal allele. As such, enhancer interactions with the *Igf2* promoter are blocked, and *H19* is expressed. On the paternal allele, enhancers are free to interact with the *Igf2* promoter and activate its expression ^9–11^. Hence, it is crucial for computational studies analyzing CTCF binding to integrate methylation data, when available, to accurately detect potential binding sites.

Mapping of CTCF-binding sites at a genome-wide scale has been performed using three broad approaches. First, computational methods that utilize CTCF PWMs and an appropriate PWM scanning tool such as the Find Individual Motif Occurrences (FIMO) ^12^ from the MEME suite or the Motif Occurrence Detection Suite of tools^13^. These methods enable genome-wide scanning of potential CTCF binding sites at a single-nucleotide resolution, with the caveat that occupancy must subsequently be inferred from orthogonal data such as chromatin state and experimental detection of CTCF protein binding. Experimental methods, such as short-read whole-genome sequencing methods, including ChIP-seq ^14,15^, CUT&RUN ^16,17^, and CUT&Tag ^18^, utilize CTCF-specific antibodies to provide direct measurements of *in vivo* CTCF occupancy. Such methodologies generate sequencing reads 50-150bp in size and hence cannot be unambiguously aligned to highly repetitive regions, such as centromeric and pericentromeric satellites. To tackle this, methods utilizing long-read sequencing have been employed. Key methods include DiMeLo-seq ^19^, DiMeLo-cito ^20^, and Fiber-Seq footprinting ^21,22^. They rely on the long reads generated by Nanopore or PacBio sequencing, and using exogenous adenine methylation, they can detect CTCF positioning in repetitive DNA sequences, with simultaneous detection of cytosine methylation at CpG sites. However, they remain technically demanding and high-priced; hence, they have not been applied for CTCF detection across many cell types. Because short-read methods fail in repetitive regions and long-read approaches are not yet scalable, computational scanning of a complete reference sequence offers a natural starting point for the systematic genome-wide characterization of CTCF binding.

Most previous human CTCF studies have been performed on the GRCh38 reference genome assembly, which leaves many repetitive regions, including centromeric and pericentromeric satellite DNA (CenSat), ribosomal DNA (rDNA) arrays on the short arms of the five acrocentric chromosomes, and several segmentally duplicated (SD) regions either missing or collapsed ^23^. The recent generation of the fully contiguous telomere-to-telomere (T2T) assembly now makes it possible to study such repetitive elements. The T2T-CHM13v2.0 assembly, provided by the T2T Consortium, originates from the uniformly homozygous CHM13hTERT cell line. It closes these gaps, introduces the remaining 8% of the human genome, and provides the first gapless representation of all 22 autosomes and the X chromosome, complemented by the HG002-derived Y chromosome ^24,25^. In parallel, the T2T Primates project has released the T2T assemblies for six non-human primates ^26^. These new assemblies enable us to characterize CTCF binding patterns in previously inaccessible regions and to compare findings for both within-primate and human-to-primate variation.

Here, we leverage these T2T assemblies to generate a comprehensive catalog of potential CTCF-binding sites. Scanning the T2T-CHM13v2.0 assembly with all three JASPAR CTCF PWMs, we discovered 39,797-56,766 sites depending on the motif, of which 12.5-15.3% could not be recovered from GRCh38. Consistent with this, 984 of the 3,604 genes newly resolved in T2T carry at least one site. We calculate CTCF density within acrocentric short arms, CenSat arrays, and SD regions. Against a GC-matched null, CTCF motifs are enriched in the α-satellite (αSat) active higher-order repeats (HORs) of chromosomes 18, 20, and X; rDNA arrays; and gamma satellites (γSats), where they present a 4.4-fold enrichment over the genome-wide average. We examine whether the similarity of acrocentric short arms leads to the identification of homologous CTCF motifs, with pseudo-homologous regions (PHRs) arising as the predominant target for CTCF inter-chromosomal interactions.

Finally, we test if our results extend to non-human primates and analyze the conservation of potential CTCF-binding sites at a base-pair resolution. Enrichment of γSat and rDNA arrays predates the divergence of great apes and gibbons. Individual CTCF sites turn over rapidly, and 56-90% of sites in the lineage-specific subterminal heterochromatin spacers overlap ancient L1 and L2 elements. We believe that the resulting resource provides one of the most complete landscapes of where CTCF can potentially bind in the human genome and can promote future experimental work.

## Results

### Genome-wide screening for potential CTCF-binding sites in the telomere-to-telomere human genome

To systematically map CTCF-binding sites across the gapless T2T-CHM13v2.0 human genome assembly, we employed FIMO with the three available PWMs for CTCF. We identified 39,797, 55,499, and 56,766 CTCF sites (FIMO *p* < 10^-6^) for MA0139.2, MA1929.2, and MA1930.2, respectively (**Supplementary Table 1**). Unless otherwise stated, all subsequent analyses are reported in this order: MA0139.2, MA1929.2, and MA1930.2. In our analyses, CTCF site density was expressed as the number of sites divided by the region’s length in Mb. When examining the per-chromosome CTCF site density, we observed that chromosome 19 had the highest density with 114 sites/Mb, and chromosome Y had the lowest with only 17 sites/Mb (**Figure 1a**). Notably, for chromosomes 18, 20, and X, the highest concentration of CTCF-binding sites was localized within the CenSat regions, suggesting a potential structural or regulatory role of CTCF at these loci (**Figure 1a**).

**Figure 1:**
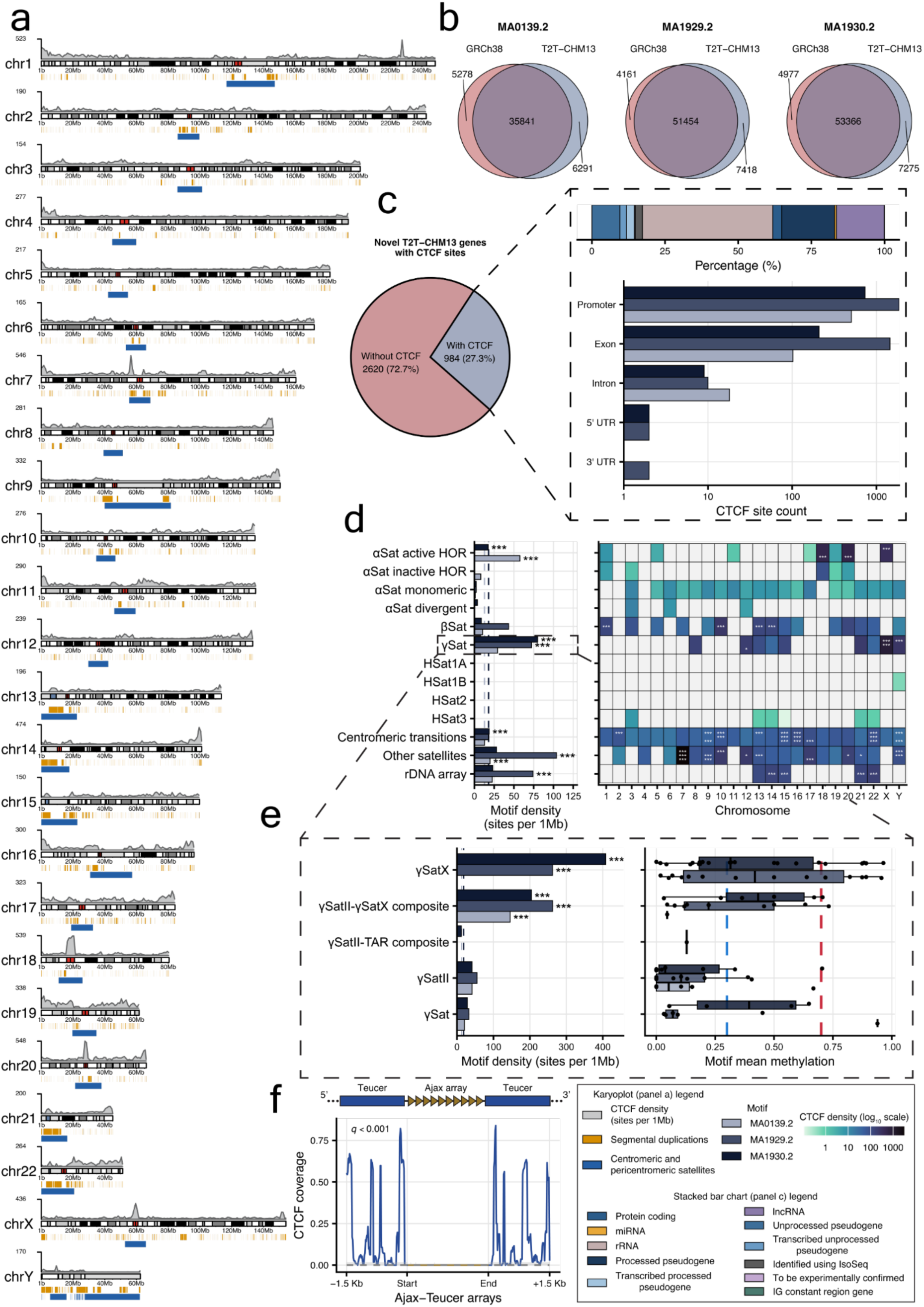
Landscape of potential CTCF-binding sites across the complete human genome assembly. **a.** Karyoplot depicting combined CTCF-binding site density across chromosomes. SDs (orange) and CenSat satellites (blue) are also depicted. **b.** Venn diagrams displaying the number of shared and assembly-specific CTCF sites between GRCh38.p14 and T2T-CHM13v2.0. **c. Left.** Pie chart depicting the proportion of T2T-CHM13v2.0 novel genes harboring CTCF-binding sites. **Right.** Stacked bar chart showing the distribution of gene categories among CTCF-bearing genes, followed by a grouped bar chart showing the number of sites across different genic compartments. **d. Left.** Bar plot showing per-motif CTCF density across CenSat types. **Right.** Heatmap depicting per-chromosome density across CenSat types. Density is plotted on a log_10_ scale. Gray squares indicate that zero potential CTCF sites were identified. **e. Left.** Bar plot displaying CTCF density per γSat subfamily. **Right.** Boxplots with jittered points showing the mean CpG methylation for each γSat-overlapping CTCF motif. The blue dashed line (methylation = 0.3) depicts hypomethylated regions, and the red dashed line (methylation = 0.7) depicts hypermethylated regions ^33^. **f.** Metaprofile plot centered on Ajax arrays, showcasing CTCF site coverage (in 15 bp bins), within arrays and their 1.5 Kb flanking regions. The gray dashed line depicts the null distribution mean, and the shaded band shows the 2.5th and 97.5th percentiles of the null. In panels d and e, the dashed lines represent the genome-wide CTCF density for each motif, and stars indicate *q* values. *: *q* < 0.05, **: *q* < 0.01, ***: *q* < 0.001. IG: immunoglobulin

To assess how many of the T2T-CHM13v2.0-identified sites represent novel findings enabled by long-read sequencing, we compared them against the GRCh38.p14 genome assembly. Using the same three CTCF PWMs, we scanned the GRCh38.p14 human genome, and we detected 41,119, 55,615, and 58,343 CTCF sites (FIMO *p* < 10^-6^), respectively. Using a progressive Cactus alignment, we lifted over the T2T-CHM13v2.0 CTCF-binding sites for all three motifs to GRCh38.p14 coordinates. The majority mapped successfully (**Figure 1b**), while 6,291 (15.3%), 7,418 (13.3%), and 7,275 (12.5%) were unique to T2T-CHM13v2.0, respectively. 5,278, 4,161, and 4,977 sites were identified exclusively in GRCh38.p14, respectively (**Figure 1b**). These CTCF-binding site differences likely correspond to previously unresolved genomic regions, genomic variation, and fixes made to issues present in GRCh38.p14, consistent with the known improvements introduced by the T2T assembly.

### Genes exclusive to the telomere-to-telomere human assembly harbor CTCF-binding sites

The T2T assembly identified genes absent from the GRCh38.p14 release of the human genome ^24^. These genes represent corrections to previous assembly errors and the detection of novel genes in previously unresolved repetitive regions. We examined their overlap with our genome-wide CTCF binding predictions to determine whether those genes contain potential regulatory elements. Out of the 3,604 new genes, 984 (27.3%) contained at least one CTCF-binding site (**Figure 1c**). They spanned multiple gene categories, with the most prominent ones being rRNA genes (*n* = 438, 44.5%), processed pseudogenes (*n* = 178, 18.1%), and lncRNA genes (*n* = 161, 16.4%) (**Figure 1c**). The predominance of rRNA genes within this set is consistent with the known role of CTCF in organizing rDNA repeats alongside cohesin ^27^.

Additionally, we examined the distribution of these CTCF sites across the promoters, exons, introns, 5’ and 3’ untranslated regions of these genes. We found that promoter and exonic regions harbored the most binding sites, with promoters having 504, 1,862, and 732 novel sites, respectively (**Figure 1c**). Out of these, we highlighted *G0001310*, *G0002160*, and *G0002132*, whose closest GENCODE gene names are *CCL3L1*, *DUX4*, and *DUX4L1*, respectively, and have been characterized as medically relevant by the Genome In A Bottle consortium ^28^. We identified potential CTCF sites in the promoters of *G0002160* and *G0002132,* suggesting that these loci may be subject to CTCF-mediated transcriptional regulation.

### Differential enrichment of CTCF-binding sites across rDNA loci, centromeric, and pericentromeric satellite classes

Prompted by the finding that potential CTCF-binding sites are present in the CenSat arrays of chromosomes 18, 20, and X, we analyzed the distribution of these sites across the genome-wide set of CenSat annotations. We obtained CenSat annotations for the T2T-CHM13v2.0 assembly ^29^, and using a GC-stratified permutation-based enrichment analysis, we identified statistically significant associations between CTCF and specific CenSat classes. αSat active HORs are enriched for two CTCF motifs (*q* < 0.001), with this enrichment being constrained only to chromosomes 18, 20, and X (*q* < 0.001) (**Figure 1d)**. However, long-read DiMeLo-seq analysis for CTCF binding in these αSat-active HORs did not produce a significant signal.

When analyzing the other CenSat classes, we observe that CTCF motifs are depleted in the remaining αSat types and the human satellites 1A, 1B, 2, and 3 (**Figure 1d**). We also detected significant enrichment of CTCF-binding sites in rDNA arrays (**Figure 1d**). Specifically, the MA1929.2 motif exhibited the highest enrichment with a density of 74 sites/1Mb (*q* < 0.001). This enriched CTCF density was observed in all rDNA arrays across the five acrocentric chromosomes, of which only chromosome 13 was not found to be statistically significant. CTCF binding in rDNA arrays was also captured in DiMeLo-seq data (**Supplementary Figure 1a**). Pericentromeric beta satellites (βSat) also showed significant enrichment of CTCF-binding sites; however, no enrichment was detected at the genome-wide level. Instead, chromosome-level analysis revealed significant enrichment (*q* < 0.001) on chromosomes 1, 10, 13, and 14, with densities ranging from 13 to 231 sites/1Mb (**Figure 1d**).

Apart from the resolved sequences of repetitive CenSat classes, the T2T human assembly enabled the annotation of the TELO_Comp element, a highly complex composite repeat element found on 10 chromosomes ^30^. Within TELO_Comp, a variable-length array of the 49-bp ajax satellite is bounded by a duplicated flanking sequence, named teucer. Applying the same GC-stratified permutation analysis, we found that ajax arrays were entirely devoid of CTCF sites (0 sites/1Mb), whereas the flanking teucer sequences were highly enriched with 5,023 sites/1Mb (*q* < 0.001) (**Figure 1f**). This effect is observed across all 10 ajax-bearing chromosomes, with chromosome 3 exhibiting the lowest density and chromosome 7. Notably, chromosome 7 also carries the most ajax arrays (*n* = 99). CTCF DiMeLo-seq data exhibited a modest binding signal in teucer arrays (**Supplementary Figure 1b**).

### Gamma satellite X harbors high CTCF density in a partially methylated environment

γSats are GC-rich tandemly repeated arrays consisting of ∼215 bp monomers ^29,31^. γSats were among the satellite classes with the highest CTCF motif density (**Figure 1d**). Prior work on the gapped genome proposed that CTCF binding in γSats functions as a barrier against centromeric heterochromatin spreading, a role that should be sensitive to the local methylation state. To study this association, we examined γSat CTCF and the methylation state of these sites. CTCF density reached 80 sites/1Mb for MA1930.2 in γSats, presenting a 4.4-fold increase over the genome-wide density (*q* < 0.001) (**Figure 1d**). Potential sites were detected in nine chromosomes; however, only chromosomes 12, X, and Y were statistically significant when accounting for GC-content. Chromosome X had the highest density with 862 sites/1Mb (*q* < 0.001), followed by chromosome Y (244 sites/1Mb) and chromosome 12 (181 sites/1Mb), combined across the three motifs (**Figure 1d**).

Since CTCF binding is sensitive to CpG methylation ^8^, we analyzed whether the CTCF sites within the different γSat subclasses reside in genomic regions permissive for CTCF binding. We obtained single-nucleotide genome-wide methylation levels generated using ultra-long nanopore sequencing and analyzed with Nanopolish ^32^. Sites with mean methylation < 0.3 were considered demethylated, sites with mean methylation between 0.3 and 0.7 moderately methylated, and sites with a mean methylation > 0.7 heavily methylated ^33^. Additionally, sites that resided in regions where methylation could not be calculated were excluded.

γSats are organized into three subfamilies: γSat, γSatII, and γSatX, sharing ∼60% sequence identity ^34^. Composite annotations between subfamilies also exist. γSat and γSatX were initially characterized in the pericentromeric regions of chromosomes 8 ^35^, and X ^36^, respectively. Subsequent analyses have detected γSat arrays across nearly all human chromosomes ^29^. γSatX exhibited the highest density, with a mean density of 223 sites/1Mb (*q* < 0.001; MA1929.2, MA1930.2), accompanied by a median methylation of 0.325 (**Figure 1e**). 44.45% of total sites were demethylated, 27.78% moderately methylated, and 27.78% heavily methylated. γSatII-γSatX composite annotations had a mean density of 204 sites/1Mb (*q* < 0.001 for all motifs) and a median methylation of 0.319 (**Figure 1e**). 50.00% of sites were demethylated, 37.50% moderately methylated, and 12.50% heavily methylated.

### CTCF sites concentrate in rDNA and satellite arrays in acrocentric short arms

The T2T-CHM13 was the first human genome assembly to sequence the short arms of the five acrocentric chromosomes accurately. These arms follow a similar structure, containing a high number of satellite repeats, rDNA arrays, and SDs ^24^. To resolve where potential CTCF sites reside within them, we computed CTCF density in 10 Kb windows across each short arm. This was accompanied by long-read CpG methylation ^32^, Fiber-seq accessibility ^37^, and DiMeLo-seq histone modification data ^38^ (**Figure 2a**). As stated previously, the rDNA arrays were amongst the regions with the highest density. This enrichment is accompanied by hypomethylation (0.213 on average) and local increased accessibility (**Figure 2a, b**). βSats also exhibited high density, especially the LSAU-BSAT composite satellites, averaging 102 CTCF sites per 1 Mb (**Figure 2a**) and a mean methylation value of 0.317. However, these regions were not found to be accessible for protein binding (mean accessibility equal to 1.398). On chromosome 15, the p-arm telomere-associated region (TAR) was the only TAR that exhibited high density, with 14 CTCF sites in 6,007 bp, whereas TARs present in the other short arms did not harbor any CTCF sites (**Figure 2a**).

**Figure 2:**
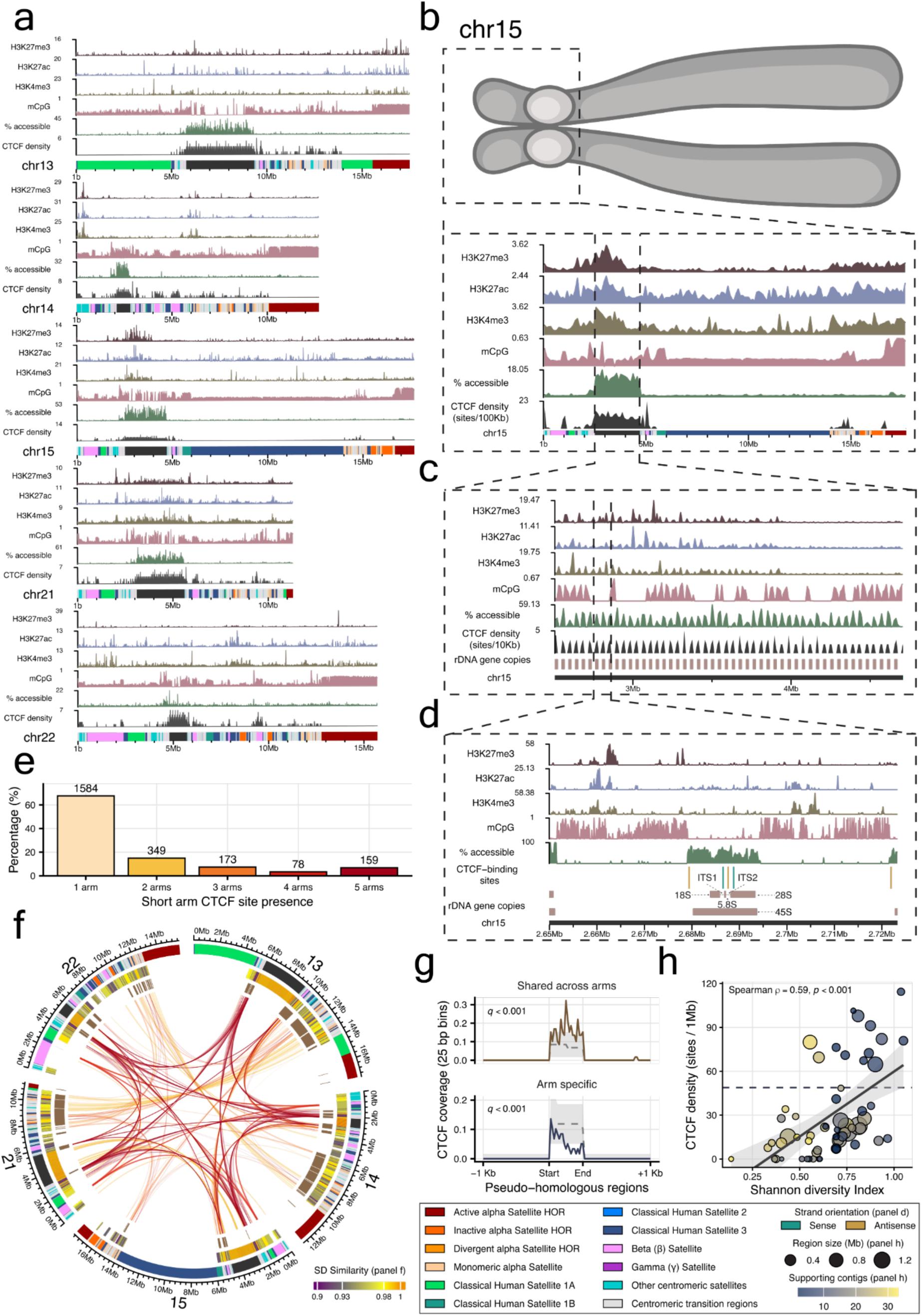
Mapping of CTCF binding in the short arms of acrocentric chromosomes. **a.** Each short arm is shown alongside the following tracks from top to bottom: H3K27me3, H3K27ac, and H3K4me3 occupancy, CpG methylation, percentage of accessible chromatin fibers, CTCF site density, and satellite annotations. All tracks are binned in 10 Kb windows to depict local densities. **b. c. d.** Multi-resolution schematic of CTCF binding in the acrocentric short arm of chromosome 15. Across resolutions, tracks follow the same order as panel a. **b.** Ideogram of chromosome 15, with the acrocentric short arm highlighted, followed by an overview of the acrocentric short arm region. **c.** Zoomed view of the rDNA array region, with bottom rectangles depicting the rDNA gene copies. **d.** A single rDNA repeat unit showing individual CTCF-binding sites (rectangles) alongside the annotated rDNA gene structure. Teal and gold rectangles indicate CTCF sites in the sense and antisense orientations, respectively. **e.** Bar plot showing the number of homologous CTCF sites that are identified across acrocentric short arms. On top of each bar, the exact number of sites is depicted. **f.** Circos plot illustrating the relationships of homologous CTCF sites between short arms. Tracks from outer to inner: CenSat; SDs; PHRs; chord links, which regions share the same site, colored by the number of chromosomes that share them. **g.** Metaprofile plot centered on PHRs, depicting CTCF site coverage within them and their 1 Kb flanks. Coverage is binned in 25 bp windows. The gray dashed line depicts the null distribution mean, and the shaded band shows the 2.5th and 97.5th percentiles of the null distribution at each bin. BH-adjusted *p* values from the permutation test are in the upper left corner. **h.** Scatter plot showing the relationship between SDI and CTCF site density across acrocentric short-arm PHRs. Each point represents one PHR; point size reflects region length and color indicates the number of supporting contigs. The dashed horizontal line marks the combined genome-wide density (sites/1Mb) across all CTCF motifs. The shaded band and line show a linear model fit (± 95% CI). Spearman’s *ρ* and *p-*value are shown in the upper left corner.

### Identification of novel CTCF-binding sites within rDNA internal transcribed spacers

Given the finding that rDNA arrays are enriched for potential CTCF-binding sites (**Figure 1c, d, Figure 2a**), we analyzed the rDNA region of chromosome 15 in detail to characterize the exact positioning of CTCF motifs within the repeat unit. High CTCF motif presence is accompanied by a local drop in methylation and an increase in accessibility (**Figure 2b**). This suggests a locally permissive epigenetic environment favoring CTCF binding within the array. Delving deeper, we observed that CTCF density, methylation, and accessibility occur in a tandemly repeated manner (**Figure 2c**). CTCF density reaches a peak within each repeat unit, with an average of 5 sites across all 50 complete rDNA units present in chromosome 15. Methylation levels are above the hypomethylation threshold between units (**Figure 2c**). Zooming into a single rDNA unit, we detect a potential CTCF site in the proximal promoter region ∼900 bp upstream of the RNA-45S pre-rRNA gene start (**Figure 2d**). The site’s methylation level is 0, and the accessibility is 75. CTCF DiMeLo-seq signal over these promoter-associated sites validated this binding pattern (**Supplementary Figure 1c**). Furthermore, we identified convergent, hypomethylated, and accessible CTCF motifs within the internal transcribed spacers (ITS1 and ITS2), consistently positioned ∼1 Kb apart (**Figure 2d**).

### Acrocentric short-arm CTCF sites are largely arm-specific, with cross-arm sites identified predominantly in centromeric transition regions

The short arms of the acrocentrics possess a high sequence similarity, with a median identity of 98.7%, measured in 5 Kb windows ^24^. This homology promotes recombination between non-homologous chromosomes ^39,40^. Due to this similarity, a CTCF site detected at a given location on one arm should also appear at the corresponding location on the other. To test this, we projected CTCF sites onto a reference-free genome-variation graph generated from the short-arm sequences of the five acrocentrics. 2,343 short-arm CTCF sites were successfully projected onto the graph. Out of these, 1,584 were arm-specific; 349, 173, 78, and 159 were identified in 2, 3, 4, and 5 arms, respectively (**Figure 2e**). Arm-specific sites were predominantly positioned within rDNA arrays (65.05%) and βSats (15.27%), whereas sites identified across arms were detected mostly in centromeric transition regions (45.85%) (**Figure 2f**).

### Shared CTCF sites are enriched in the recombining pseudo-homologous regions of the acrocentric short arms

Pseudo-homologous regions (PHRs) are stretches of near-identical sequence shared across the short arms of the acrocentric chromosomes. They exhibit population-level signatures of ongoing recombination exchange between non-homologous chromosomes ^39^. They are analogous to the pseudoautosomal regions found in the X and Y chromosomes, where otherwise non-homologous chromosomes pair and recombine ^39,41^. An initial analysis found that 47.30% of CTCF sites shared across short arms fall within PHRs (**Figure 2f**). To test whether this enrichment is statistically significant and specific to shared sites, we performed a GC-stratified permutation-based analysis. Examining their distribution within PHRs and their 1 Kb flanks, we found that homologous CTCF sites, found in more than 1 arm, occur 1.7-fold more often in PHRs (*q* < 0.001) than expected by chance or sequence composition alone (**Figure 2g**). By contrast, arm-specific sites were depleted in PHRs, occurring 1.58-fold less often than expected by the same GC-stratified null (*q* < 0.001) (**Figure 2g**).

Next, we tested whether there was a correlation between a PHR’s CTCF density and its Shannon diversity index (SDI). The SDI is a metric used to quantify how many different acrocentric chromosomes a region’s sequence resembles across the population. A value near zero means a region matches a single chromosome, while higher values indicate it is a mosaic of several ^39^. We identified a statistically significant correlation between CTCF density and SDI (Spearman’s *ρ* = 0.59, *p* < 0.001) (**Figure 2h**). Fitting a Negative Binomial model confirmed that this relationship remained positive after adjusting for GC content and region length, though the association was weaker than the bivariate correlation (*β* = 1.29, *p* = 0.025) (**Figure 2h**). These results indicate that potential CTCF sites accumulate in proportion to how many acrocentric chromosomes a region assembles and are concentrated in the most mosaic, recombination-prone PHR sequences.

### CTCF enrichment within γSat and rDNA arrays predates the divergence of great apes and gibbons

Since we observed that CenSats harbor significant patterns of CTCF-binding sites in the human genome assembly, we then asked whether these also expand to primates. To address this, we applied the same methodology across six T2T non-human primate assemblies and their corresponding CenSat annotations. The non-human primates analyzed were *Gorilla gorilla gorilla*, *Pan paniscus*, *Pan troglodytes*, *Pongo abelii*, *Pongo pygmaeus*, and *Symphalangus syndactylus* (**Supplementary Table 2**).

Across the seven primate species, we observed that γSat was substantially enriched, with a mean density of 1,190 sites/Mb (*q* < 0.001 for multiple motifs across species). rDNA arrays also exhibited species-wide enrichment with a mean density of 235 sites/Mb (*q* < 0.001 for multiple motifs across species) (**Figure 3a**). These patterns suggest that the association of CTCF motifs with γSats and rDNA arrays predates the divergence of great apes and gibbons. Centromeric transition regions were also found enriched across species with a mean density of 65 sites/1Mb (*q* < 0.001) (**Figure 3a**). In humans, CTCF binding in centromeric transition regions is also supported by orthogonal DiMeLo-seq data (**Supplementary Figure 1d**).

**Figure 3:**
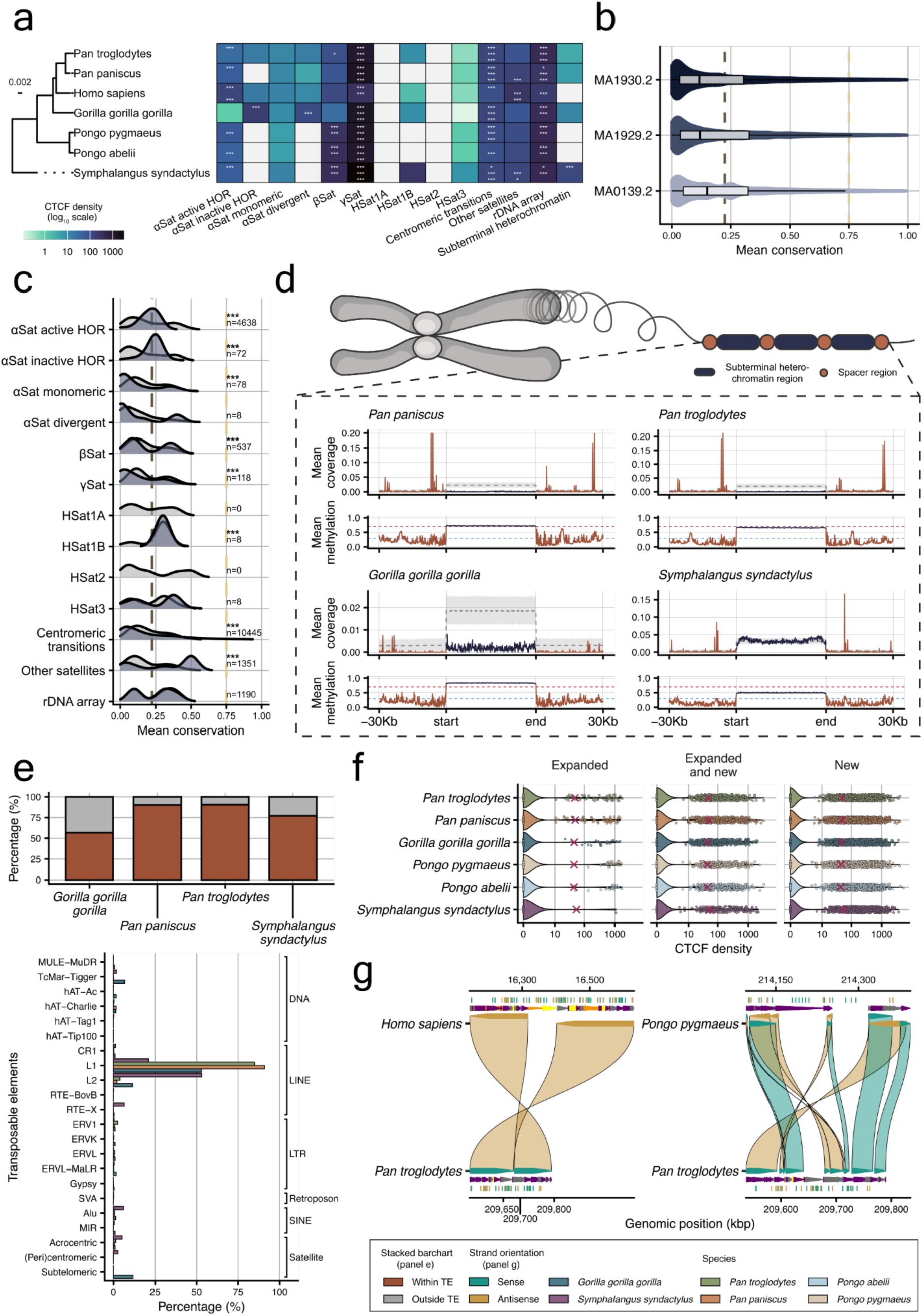
Comparative analysis of CTCF-binding sites across primate telomere-to-telomere genomic assemblies. **a. Left.** Phylogenetic tree of *Homo sapiens*, *Gorilla gorilla gorilla*, *Pan paniscus*, *Pan troglodytes*, *Pongo abelii*, *Pongo pygmaeus*, and *Symphalangus syndactylus*. **Right.** Heatmap depicting CenSat CTCF density (sites/1Mb) across primate species. Density is plotted on a log_10_ scale. Gray squares indicate that zero potential CTCF sites were identified. Stars indicate Benjamini-Hochberg adjusted *p-*values. *: *q* < 0.05, **: *q* < 0.01, ***: *q* < 0.001. **b.** Beeswarm plot depicting the distribution of conservation scores of CTCF sites across the three motif types. Each beeswarm plot is overlaid by a boxplot showing median values and interquartile ranges. **c.** Ridgeline plot showing CTCF site conservation across the different CenSat categories. For each category, *n* represents the number of sites overlapping it. In panels **b** and **c**, the brown dashed line represents the mean genome-wide conservation score (score = 0.225), and the gold line represents the threshold used to determine highly conserved CTCF sites (score = 0.75). **d.** Illustrative schematic showing the tandemly repeated structure of subterminal heterochromatin regions, followed by metaprofile plots portraying the mean CTCF site coverage and mean site methylation across all regions and their 30 Kb flanking windows split into 100bp bins. The gray dashed line depicts the null distribution mean, and the shaded band shows the 2.5th and 97.5th percentiles of the null distribution at each bin. The colored dashed lines in the methylation tracks indicate hypomethylation (methylation=0.3; blue line) and hypermethylation (methylation=0.7; red line) thresholds. **e. Top panel.** Stacked bar chart showing the fraction of CTCF sites in subterminal heterochromatin spacers that overlap transposable elements. **Bottom panel.** Grouped bar chart depicting the distribution of these overlapping sites across transposable element subclasses. **f.** Jittered point violin plots showing CTCF density across lineage-specific segmentally duplicated regions. Each point represents a unique region, while the “x” mark represents the genome-wide CTCF density for each species. The x-axis is log-scaled. **g.** Alignment view of chr1: 209,537,154-209,792,028 of *Pan troglodytes*, aligned to *Homo sapiens* (left) and to *Pongo pygmaeus*. For each alignment, the individual CTCF motifs and SDs are also depicted.

We also detected species-specific patterns. *Gorilla gorilla gorilla* had the lowest CTCF density (0.3 sites/1Mb) in αSat active HORs; meanwhile, the rest of the primates had a mean density of 32 sites/1Mb (*q* < 0.001 across species). Interestingly, *Gorilla gorilla gorilla* was the only species with a substantial CTCF density in αSat inactive and divergent HORs, with 107 sites/1Mb and 55 sites/1 Mb, respectively (*q* < 0.001 for both). This pattern amounts to a 2.3x and 1.2x higher density than the genome-wide average (**Figure 3a)**.

### Conservation and divergence of CTCF sites across primate centromeric satellites

Next, we analyzed the conservation of CTCF sites at base-pair resolution, first genome-wide and then within each CenSat class. We obtained PhastCons conservation scores for the seven primates, and we assigned a score to each potential CTCF site based on the mean per-base score across the motif span. The majority of CTCF sites were species-specific, with the conservation score appearing the most being 0.024, 0.024, and 0.022 across motifs, roughly an order of magnitude less conserved than the genome-wide average of 0.225. However, we were also able to identify 1,385, 2,116, and 2,056 highly conserved CTCF sites, respectively (**Figure 3b**).

When examining the CTCF site conservation across the different CenSat categories, we found that most sites within aSat active and inactive HORs approach the genome-wide conservation score (mode equal to 0.25 for both categories), and their score distribution differs significantly from that of the surrounding region, with modes equal to 0.1 and 0.06 for active and inactive αSats, respectively (*q* < 0.001; Kolmogorov-Smirnov test) (**Figure 3c**). The distribution of conservation scores of CTCF sites within βSats and γSats was also found to be statistically different from that of the surrounding regions (*q* < 0.001; Kolmogorov-Smirnov test), with βSat CTCF sites following a bimodal distribution, whereas γSat CTCF sites were primarily found to be highly diverged, with the score appearing the most being 0.02, 4.5-fold lower than the γSat mode of 0.1 and 8.9-fold lower than the genome-wide average. Centromeric transitions proved to be an interesting example, since it was the only class containing highly conserved CTCF sites (n=296 sites), with this effect being observed across chromosomes (**Supplementary Figure 2**).

### CTCF sites are depleted in pCht subterminal caps but enriched in spacers and αSat caps of non-human primates

The complete sequencing of non-human primates also revealed the sequence of the subterminal heterochromatin caps, large blocks of tandemly repeated DNA ^26,42^. These satellites are conspicuously absent from the human lineage and are present only in *Gorilla gorilla gorilla, Pan paniscus, Pan troglodytes,* and *Symphalangus syndactylus*. They account for a substantial portion of primate genomes, with coverage ranging from 4% in *Pan troglodytes* to 10.1% in *Symphalangus syndactylus ^26^*. In *Gorilla gorilla gorilla* and the *Pan* lineage, the subterminal heterochromatin regions consist of a 32 bp AT-rich satellite sequence, named pCht, whereas in *Symphalangus syndactylus*, the monomer sequence is a 171 bp αSat ^43,44^; in both cases, spacer sequences are present between satellite arrays (**Figure 3d**).

To determine whether CTCF-binding sites are present within these regions, we used the FIMO-derived sites across all three motifs for the four non-human primates harboring these heterochromatin caps. We also examined the methylation state of both subterminal caps and spacer regions, using Nanopore long-read methylation calls. Both site coverage and mean CpG methylation were calculated across each region and its 30 Kb flanks, partitioned into 100 bp bins (**Figure 3d**). Species of the *Pan* lineage exhibited similar enrichment patterns, with CTCF sites depleted within the heterochromatin caps and spacer regions harboring peaks. These peaks were observed ∼6 Kb and ∼26 Kb upstream and downstream, respectively, accompanied by a mean methylation of 0.336 and 0.451 across the two species (**Figure 3d**).

*Gorilla gorilla gorilla* had the lowest CTCF coverage of the four species. Despite sharing the same pCht monomer with the *Pan* species, coverage was an order of magnitude lower, while the methylation profile was highly similar, with hypermethylation (scores > 0.7) within the caps and hypomethylation (scores < 0.3) in the spacers. This difference in potential CTCF binding can be partially explained by the fact that these spacer sequences have different evolutionary timing (∼7.7 and ∼5.0 million years in *Pan* and *Gorilla gorilla gorilla*, respectively). Additionally, they possess different ancestral origins (intronic region of *MALD1* for *Gorilla gorilla gorilla* and a subsequence of *PGM5* for *Pan* species) ^42^. *Symphalangus syndactylus* was the only species that displayed CTCF enrichment in both subterminal and spacer regions. The subterminal regions exhibited a density of 62 sites/1Mb (*q* < 0.001 for MA1930.2) (**Figure 3a**), while the highest peak occurred ∼8 Kb downstream of the subterminal heterochromatin cap (**Figure 3d**).

### Ancient LINE elements are the predominant source of CTCF sites within subterminal heterochromatin spacer regions

After observing high coverage of CTCF sites in the spacer regions, we analyzed their overlap with transposable elements, since the spacer regions constitute pockets of euchromatin DNA with a high content of transposable elements ^42^. For each non-human primate assembly with subterminal heterochromatin regions, we used transposable element annotation obtained from RepeatMasker ^45^, alongside the Dfam database (v3.6) ^46^. Initially, we examined the fraction of spacer CTCF sites overlapping such elements. We observed that across all species, more than 50% of sites overlapped, ranging from 56.72% in *Gorilla gorilla gorilla* to 90.46% in *Pan troglodytes* (**Figure 3e**). When analyzing transposable element distribution among these overlapping sites, we found that the overlap was predominantly driven by long interspersed elements (LINEs) (**Figure 3e**).

Specifically, L1 elements accounted for 90.94%, 84.90%, 52.86%, and 21.22% of CTCF sites in spacers of *Pan paniscus*, *Pan troglodytes*, *Gorilla gorilla gorilla*, and *Symphalangus syndactylus*, respectively. For *Symphalangus syndactylus*, L2 elements were more predominant, accounting for 53.07% of sites (**Figure 3e**). L1ME4b was the element appearing the most in both *Pan* species, with coverages of 41.29% and 38.98%, in *Pan paniscus* and *Pan troglodytes*, respectively, whereas L2d2 (37.02%) and L1MC (12.46%) were the most abundant elements in *Gorilla gorilla gorilla* and *Symphalangus syndactylus* spacer CTCF sites, respectively (**Supplementary Table 3**). These findings align well with the established model, in which transposable elements act as an evolutionary source of CTCF sites in mammalian genomes ^47,48^. L1 elements have been found to carry CTCF sites within their sequences and can contribute to species-specific CTCF binding ^49,50^. Whole-genome defragmentation analysis performed by Giordano *et al.* has established L2 elements, L1MC, and L1ME4 to be amongst the chronologically oldest LINE elements ^51^, suggesting that spacer CTCF sites derive from ancestrally embedded motifs rather than recent retrotransposition.

### Primate lineage-specific segmental duplications can lead to *de novo* CTCF site acquisition

Due to the use of long-read sequencing, T2T non-human primate genomes have enabled the identification of previously unknown SD regions ^26^. Chaining SDs within 100 Kb and comparing the homologous loci across lineages allowed certain regions to be classified as lineage-specific ^26^. These regions were organized into three categories: *expanded*, showing a lineage-specific increase in copy number; *new*, arising from lineage-specific sequence change; and *expanded and new*, exhibiting both. We computed potential CTCF site density within each region to examine how lineage-specific duplications affect CTCF binding sites.

CTCF sites were detected in all lineage-specific SD categories across the six non-human primates. In all categories, the majority of regions (77.2 - 81.5%) exhibited modest densities, which were below each species’ genome-wide density (**Figure 3f**). However, each one contained certain regions with highly elevated density, reaching up to 5,240 sites/1Mb for *Pongo abelii* in the category *new*. *New* and *expanded and new* SDs possessed an average of 21.4% and 22.4% of regions with density greater than the genome-wide average, respectively, whereas the *expanded* category showcased only a 10.2% (**Figure 3f**).

To resolve how these sites are arranged within a single lineage-specific duplication, we highlight here one example of a Homininae-specific SD, identified in chromosome 1 of *Pan troglodytes* (region: 209,537,154-209,792,028). We aligned this sequence to the T2T assemblies of the other primates and selected the alignments of *Homo sapiens* and *Pongo pygmaeus* to highlight (**Figure 3g**). This region exhibited a density of 196 sites/1Mb and was categorized as *new*. We observed that this SD region mapped to a single *Homo sapiens* locus on chromosome 1 as two inverted blocks spanning ∼172 Kb and ∼228 Kb, separated by a ∼85 Kb human-specific insertion (**Figure 3g**). In *Pongo pygmaeus*, the SD collapses onto a single ∼138 Kb locus, with distinct *Pan troglodytes* intervals aligning to overlapping coordinates in opposite directions (**Figure 3g**). Of the 50 potential CTCF sites in the Homininae-specific SD, 86% had a corresponding site in the *Homo sapiens* assembly, whereas only 14% did in *Pongo pygmaeus* (**Figure 3g**), despite ∼77% of the locus being aligned between the two species. This finding suggests that most CTCF sites in this region are not inherited from the ancestral locus, implying the sites were introduced in the Homininae subfamily together with the duplication.

## Discussion

The completion of the human genome in 2022 by Nurk *et al*. ^24^ resolved roughly 8% of the sequence, consisting primarily of repetitive, heterochromatic regions, which prior assemblies left collapsed or absent. Hence, CTCF’s role within these regions had proven hard to study until recently. Here, we scanned the T2T-CHM13v2.0 assembly, together with six non-human primate T2T assemblies, for potential CTCF-binding sites and found that these newly accessible compartments are not depleted of CTCF motifs. CenSat regions, rDNA arrays, acrocentric short arms, and the lineage-specific subterminal heterochromatin caps of great apes and gibbons are enriched for CTCF sites when tested against a null distribution of 10,000 permutations, all accounting for GC content. Our findings suggest that CTCF’s reach into the repetitive genome has been underestimated in the past, potentially due to assembly incompleteness and not due to biology.

Previous work in mouse erythroleukemia cell systems, using synthetic repetitive DNA vectors, had shown that CTCF binding in γSats allows for a permissive chromatin state ^31^. With the resolved human γSat sequence, we show that this pattern is consistent in humans, suggesting CTCF as an insulator blocking heterochromatin spreading from the active centromere core to the euchromatic flanks. Additionally, we show potential CTCF binding within the αSat active HORs, responsible for kinetochore nucleation and segregation of sister chromatids during mitosis ^52^. CTCF degradation in HCT116 cells has been shown to lead to mitotic failure, with increased intercentromere distances and a more disorganized metaphase plate ^53^. Such phenotypes were observed approximately three days after CTCF degradation and mirror the effects of partial cohesin loss, suggesting that CTCF maintains rather than establishes the centromeric chromatin structure. Whether this involves CENP-E remains contested: CTCF has been reported to recruit CENP-E to the pericentromeric/centromeric regions via unusual CTCF binding sites ^54^, but acute CTCF degradation leaves CENP-E kinetochore levels intact ^53^. Such discrepancies possibly reflect differences in depletion duration.

rDNA arrays also arose as a prominent CTCF target, with the majority of CHM13 novel genes carrying CTCF sites being rDNA-associated genes. The rDNA promoter sites we report align with the previously identified sites present in the H42.1 promoter ^55^. Henikoff and Henikoff reported that CTCF binding within the rDNA spacer promoter flanks the miR--1275/miR--6724 transcription unit within a <400 bp window, with CTCF degradation reducing PRO-seq signal at this locus ^56^. Additionally, we identify binding sites in both internal transcribed spacers of rDNA units, which, to our current knowledge, have not been reported in the past.

Beyond individual satellite classes, our short-arm analysis suggests the potential implication of CTCF in the interchromosomal exchange between acrocentric chromosomes. CTCF sites were 1.7-fold enriched in the near-identical, actively recombining PHRs. These regions behave as an autosomal analogue of the pseudoautosomal regions found on the X and Y chromosomes ^39^. CTCF density in PHRs scales with the degree of sequence mosaicism across chromosomes. PHRs also hold clinical relevance, since they act as the substrate for Robertsonian translocations ^57^, the most common balanced chromosomal rearrangement in humans ^58^. Because the acrocentric short arms are drawn into physical proximity within the nucleolus, an architectural factor, such as CTCF, positioned at the regions of shared homology, is meant to either influence or be influenced by this interchromosomal contact.

Extending our analysis to T2T non-human primate assemblies demonstrates that CTCF binding in satellite regions is not a human-specific trait. γSats and rDNA arrays were enriched in every species examined, as were centromeric transition regions. Alpha satellites behaved differently. αSat active HORs carried motifs in six of seven species but were nearly empty in *Gorilla gorilla gorilla*, which was instead the only species with substantial density in inactive and divergent HORs. Given that αSat HOR arrays undergo a nearly complete turnover, with changes idiosyncratic to each species ^59^, our results suggest that the *Gorilla gorilla gorilla* active array has diverged beyond retention of the CTCF motif, while its relic arrays preserve the ancestral state.

In primate subterminal heterochromatin caps, CTCF sites were depleted within the hypermethylated pCht arrays and concentrated in the hypomethylated spacers between them. 57-90% of spacer sites overlapped predominantly ancient L1 and L2 LINE elements. This is consistent with the established model that retrotransposon expansion seeds mammalian CTCF sites ^47,48^. The *Gorilla*-*Pan* comparison is the most interesting, since both share the same pCht arrays, with near-identical methylation levels, yet an order of magnitude difference in potential CTCF binding is observed. *Symphalangus syndactylus* was the only primate with significant CTCF coverage both within the subterminal cap and the neighbouring spacers. This can be attributed to the fact that such caps are built from αSat arrays and not pCht ^26^.

Our work comes with certain limitations. Our sites are computational predictions of sequence-encoded binding potential, not measurements of *in vitro* or *in vivo* occupancy. A motif match indicates that CTCF can bind, but these sites can be cell type-specific, cell cycle-dependent, or rendered inaccessible by the local chromatin and methylation state. While the *p* < 10^-6^ threshold has been reported to produce ∼80% true positives, that benchmark is derived from short-read ChIP-seq data and might not extend to the repetitive regions of the genome, where no comparable ground truth exists. CTCF DiMeLo-seq data used in this study as orthogonal evidence also provide limited evidence since they are derived from a single cell line and two replicates; hence might be insufficient to draw generalizable conclusions. Finally, since CTCF binding can be both cell cycle and cell type specific, the absence of DiMeLo-seq signal at a predicted site, such as the case of γSats, cannot be taken as evidence that the site is never bound. This is especially relevant in the context of PHRs, where CTCFs’ putative role requires meiotic cells so that it can be tested, and can not be validated in the GM12878 DiMeLo-seq data we possess. Our current findings establish only co-localization, not a mechanism.

Concluding, by extending CTCF mapping into the repetitive regions that earlier assemblies could not resolve, our analysis reframes them as a potentially integral component of CTCF’s functions. The patterns we describe are sequence-level hypotheses that now invite further experimental testing. Long-read CTCF footprinting across diverse cell types, and critically in meiotic and germline contexts, would establish whether PHR-associated sites are bound when their proposed recombination role would be active. Extending this analysis from a single reference to the growing set of pangenome and primate T2T assemblies would distinguish constrained CTCF sites from those that turn over rapidly. Since the methods we have to study the genome are progressing at such a fast rate, we expect the repetitive compartments to become an important aspect of the functional genome, and we offer this resource as a foundation for that effort.

## Materials and methods

### Genome assemblies

The complete human telomere-to-telomere assembly (T2T-CHM13v2.0) was used as the primary reference assembly throughout the study. T2T-CHM13v2.0 provides a gapless representation of the 22 autosomal chromosomes and the X chromosome from the CHM13hTERT cell line and the gapless sequence of the Y chromosome from the HG002 cell line. Finally, GRCh38.p14 was used as a baseline reference to quantify CTCF motif gain due to telomere-to-telomere assembly completeness. The full assembly was used, including alternative loci, fix patches, and decoy sequences.

To assess the conservation of CTCF-binding sites across non-human species, we used the T2T-level diploid assemblies of six primate species generated by the T2T Primate Consortium. For each species, we used only the primary haplotype assembly, which represents the highest quality haplotype for each chromosome and includes both sex chromosomes. Assembly details and GenBank accession numbers are displayed in **Supplementary Table 2**.

### Prediction of genome-wide CTCF-binding sites

Potential CTCF-binding sites were obtained using the FIMO ^12^ tool from the MEME suite (v5.5.9) ^60^. Prior to motif scanning, a background model of nucleotide frequencies was generated for each analyzed genome using the fasta-get-markov tool from the same suite to improve FIMO’s statistical scoring. A p-value < 10^-6^ was used as a threshold to define statistically significant sites. This choice was based on the finding by Dozmorov *et al. ^61^* that approximately 80% of hits at this cutoff represent true positive CTCF-binding sites. Since CTCF binding can be facilitated by different zinc-finger combinations, three PWMs exist, which are stored in the JASPAR database, an online resource containing more than 2,500 transcription factor PWMs ^62^. Motif MA0139.2 models the core sequence, and motifs MA1929.2 and MA1930.2 extend this core sequence to capture contacts made by the flanking zinc finger modules. Three independent FIMO runs were performed, one for each PWM.

### Genome-wide CTCF density calculation and visualization

CTCF density is defined as the number of potential CTCF-binding sites per 1Mb of the human genome, regardless of motif orientation. The genomic density was plotted for each chromosome using karyoploteR (v1.32.0) ^63^, alongside segmental duplications, centromeric, and pericentromeric satellites.

### Mapping CTCF-binding site differences between GRCh38.p14 and T2T-CHM13v2.0 reference assemblies

The Cactus 8-way primary progressive alignment (See Code and data availability section) was filtered to keep only the alignment between GRCh38.p14 and T2T-CHM13v2.0 using HALtools (v2.2.0) ^64^. T2T-CHM13v2.0 genomic coordinates of statistically significant FIMO CTCF sites were lifted over to GRCh38.p14 coordinates using halLiftover (v2.2.0) and the HAL alignment file for each CTCF motif. After halLiftover, regions shorter than 15bp (the length of the MA0139.2 motif) were discarded as degenerate mappings. T2T-CHM13v2.0 regions that failed the lift-over process were characterized as T2T-CHM13v2.0-unique. Overlap between native GRCh38.p14 CTCF sites and GRCh38 CTCF sites lifted-over from T2T-CHM13v2.0 was assessed using the GenomicRanges (v1.58.0) ^65^ package, requiring the native GRCh38.p14 site to be fully contained within a lifted-over site. Native GRCh38.p14 CTCF sites with no lifted-over counterpart were characterized as GRCh38.p14-unique.

### Detection of overlaps between CTCF sites and regions of interest

Overlaps between CTCF-binding sites and regions of interest were identified using the Bioconductor GenomicRanges package (v1.58.0) ^65^. Overlaps were called at a minimum of 1bp overlap, regardless of strand orientation.

### Permutation-based enrichment analysis of CTCF motifs across genomic regions

To assess whether the increase in CTCF density within a specific genomic region was statistically significant, we performed a permutation-based overlap analysis using the Genomic Association Test (GAT) tool (v1.3.6) ^66^. Initially, the genome was split into 100 Kb non-overlapping bins using a sliding window approach. For each bin, we calculated the GC content (number of G and C nucleotides divided by the bin’s length), and we stratified bins into six isochore classes (L1, L2, M1, M2, H1, and H2) by partitioning the GC content distribution into six tiles. This way, genomic bins with similar GC content are assigned to the same category. For each combination of CTCF motif and genomic annotation of interest, we performed 10,000 permutations, in which motifs were randomly shuffled within regions belonging only to the same isochore, while allowing for potential overlaps. During each permutation, we recalculated the number of overlaps with the region of interest to generate a null hypothesis. This enabled us to create a null hypothesis that accounts for both the random distribution of CTCF sites and the genomic composition. All overlap tests were performed in a strand-agnostic manner. Statistically significant enrichments were considered only those with a Benjamini-Hochberg adjusted p-value less than 0.05.

### Identification of CTCF binding from DiMeLo-seq data

Raw Oxford Nanopore fast5 data were basecalled using Guppy (v6.5.7) ^67^ using the res_dna_r941_min_modbases-all-context_v001 Rerio model ^68^, which calls base modifications at all sequence contexts. Basecalling was performed without a methylation threshold to retain all modification calls for downstream filtering. Output BAM files for each sequencing run (2 replicates) were merged with SAMtools (v1.21.0) ^69^ and converted to FASTQ format with the MM/ML modification tags preserved. Meryl (v1.4.1) ^70^ was used to count repetitive 15-mers, with a distinct frequency greater than 0.9998, and these were supplied to Winnowmap (v2.0.3) ^71^ for read alignment to the T2T-CHM13v2.0 assembly. Aligned reads were coordinate-sorted and merged into a single BAM file. Modkit (v0.2.4) ^72^ was implemented to update the base modification tags to the standard ambiguous-mode format using the updated-tags command. Per-chromosome adenine (tag: A,0) and CpG (tag: CG,0) methylation pileups were calculated using the dimelo-toolkit software (v0.2.1) ^73^. Modkit’s adaptive threshold determination was used to exclude low-quality base modifications, and the final pileup BED files were filtered to keep only 6mA calls and were converted to bigWig format using the bedGraphToBigWig (v2.10.0) tool ^74^.

### Histone modification profiling from DiMeLo-seq data

H3K27ac, H3K27me3, and H3K4me3 modification data were processed following the same pipeline as described for CTCF binding identification, without the basecalling and read alignment steps, since they were obtained in pre-processed BAM format with base modification tags already present from Maslan *et al*. ^38^.

### Identification of shared CTCF sites between acrocentric chromosome short arms using a genome graph approach

Short arm genomic sequences (from the chromosome start up until the centromere’s alpha satellite active higher-order repeat, as defined by the CenSat annotations) from the five acrocentric chromosomes were extracted using the BEDTools getfasta (v2.31.1) ^75^ command. To jointly represent and compare their sequence variation, we constructed a genome graph from these sequences using the PanGenome Graph Builder (v0.7.4) ^76^ algorithm. This enabled us to create a reference-free sequence alignment to examine sequence similarity between any pair of short arm sequences. For the construction, we used a segment length of 25 Kb and a minimum mapping identity of 95%. Matches with lengths less than 100 bp were filtered during graph induction. The output genome graph was sorted using ODGI (v0.9.4) ^77^, following a path-guided stochastic gradient descent sort (ODGI flag: -p Ygs). FIMO CTCF sites previously identified within the short arm sequences were projected onto the genome graph coordinate system using the ODGI position command. We lifted each site’s coordinates from its reference chromosome onto the corresponding positions in all other acrocentric short arms. Sites were considered shared between short arms if the lifted interval fell entirely within an independently identified CTCF motif on the target arm. CTCF site sharing was visualized using the circlize (v0.4.18) R package ^78^.

### Enrichment analysis of CTCF sites in PHRs of acrocentric chromosomes

To examine whether short-arm PHRs were enriched for CTCF sites, we implemented a similar framework to the one described in the section “Permutation-based enrichment analysis of CTCF motifs across genomic regions”. The permutation analysis was performed for both sites that were unique to one short arm and for those that were shared across arms. We ran 10,000 permutations for each category. To characterize the distribution of CTCF sites within PHRs, we used the EnrichedHeatmap (v1.36.0) R package ^79^. Metaprofile plots were generated centered on PHRs with 1 Kb flanking extensions, divided into 25 bp bins. For each bin, CTCF site coverage was calculated and averaged across all PHRs. Similar methodology was applied for each one of the 10,000 permutations, and the mean signal was plotted alongside the 2.5th and 97.5th percentiles of the null distribution at each bin.

### Correlation between CTCF density and SDI

CTCF site density was calculated for each PHR, similarly to the Genome-wide CTCF density calculation and visualization section. To assess whether CTCF density varies with sequence homology between PHRs, we tested its correlation with the SDI. This index is a positional homology entropy metric that reflects the degree to which a genomic region is shared across multiple acrocentric chromosomes ^39^. Spearman correlation was used to quantify this relationship, and a Negative Binomial model was fitted using CTCF counts as the response variable, with SDI and GC content as predictors. We also included the log-transformed PHR width as an offset to account for differences in region length.

### Conservation analysis of potential CTCF-binding sites across telomere-to-telomere primate genome assemblies

To evaluate the conservation of the detected CTCF sites across the complete genomes of primates, we obtained PhastCons (PHAST v1.5.0) ^80^ per-base conservation scores from the work of Yoo *et al.* ^26^ In brief, multiple sequence alignments between the T2T-CHM13v2.0 (defined as the reference assembly) and the six primate genomes (**Supplementary Table 2**) were generated using wfmash all-to-all (v0.24.2) ^81^, and the generated PAF files (one with each assembly as a target/reference) were filtered with wgatools (v1.1.0) ^82^ to keep alignments with length greater than 10 Mb. For the conservation score estimation, a fourfold degenerate site model was generated using entire alignments for all chromosomes except for chromosome 2, which was split into 1 Mb chunks. This model was applied genome-wide, and the --target-coverage and --expected-length parameters of the PhastCons HMM were optimized using a grid search approach ^83^. This approach resulted in a mean genome-wide conservation score of 0.225. To assign a conservation score to each CTCF site identified in the T2T-CHM13v2.0 assembly, we calculated the mean per-nucleotide score over the motif’s length. We defined highly conserved CTCF sites as those with a mean score greater than 0.75.

### Analysis of CTCF site positioning within primate subterminal heterochromatin regions and associated spacer sequences

Using the same approach as described in the Enrichment analysis of CTCF sites in PHRs of acrocentric chromosomes section, we examined the distribution of CTCF sites across the subterminal heterochromatin regions of *Gorilla gorilla gorilla*, *Pan paniscus*, *Pan troglodytes*, and *Symphalangus syndactylus*. We employed the EnrichedHeatmap (v1.36.0) R package ^79^, generating metaprofile plots centered on subterminal heterochromatin regions with 30 Kb flanking extensions, divided into 100 bp bins. This was performed independently for both CTCF coverage and long-read methylation levels.

### Identification of transposable elements harboring CTCF sites within subterminal heterochromatin spacer sequences

Spacer regions were derived by first merging subterminal heterochromatin regions separated by less than 1 Mb, then subtracting the original subterminal heterochromatin intervals from the merged windows, as defined by Yoo *et al*. *^26^* Spacer-associated CTCF sites were then intersected with RepeatMasker-derived ^84^ transposable element annotations, allowing for one CTCF site to map to multiple elements, respecting the nested structures of transposable elements.

### Alignment of the *Pan troglodytes* lineage-specific SD region to the other primate assemblies

The complete genomic sequence of chromosome 1 from the *Pan troglodytes* assembly was aligned using minimap2 (v2.31-r1302) ^85^ with the options “-x asm20 -c --eqx -p 0.5 -N 50” to the genomes of the other primates and of *Homo sapiens*. Alignments were filtered to keep only those with a MAPQ score greater than 10, and the alignments were visualized using SVbyEye (v0.99.0) ^86^ alongside individual SD blocks and potential CTCF sites. Liftover of CTCF sites between homologous regions of the alignment was carried out using the liftRangesToAlignment command from SVbyEye. Liftover sites that also overlapped CTCF annotations in the target region (minimum 8bp overlap) were considered as homologous CTCF sites.

## Code and data availability

CTCF motif file pre-processing was performed using a custom pipeline built with Snakemake (version 7.32.4) and GNU Bash (version 5.2.32), and downstream statistical analyses were performed in R (version 4.4.2) using various base and additional packages. All scripts developed for this study are available in the following GitHub repository: https://github.com/Georgakopoulos-Soares-lab/T2T_CTCF. Scripts are published under the GNU General Public Licence v3.0, allowing researchers and interested individuals to access and use the custom scripts for their own analyses or to replicate the study’s findings.

CTCF PWMs used in this study were obtained from the JASPAR CORE database (2026 release) ^62^. PWMs MA0139.2, MA1929.2, and MA1930.2 were used across all analyses.

The T2T-CHM13v2.0 reference genome (<u>chm13v2.0.fa</u>), along with associated cytoband (<u>chm13v2.0_cytobands_allchrs.bed</u>) and CenSat annotations (<u>chm13v2.0_censat_v2.1.bed</u>), was obtained from the official T2T consortium’s AWS bucket. CHM13hTERT nanopore methylation data were obtained from the Gershman *et al.* study ^32^. The GRCh38.p14 reference genome was obtained from the <u>NCBI Genome FTP repository</u>. Raw long-read DiMeLo-seq data of CTCF binding in GM12878 cells were obtained from the work of Altemose *et al*. ^19^ under the SRA accessions SRR18623941 and SRR18623919. Pre-processed H3K27ac, H3K27me3, and H3K4me3 DiMeLo-seq data from GM12878 cells were obtained from Maslan *et al*. ^38^, deposited in GEO under accession GSE208125. Pre-processed Fiber-seq accessibility data from CHM13hTERT cells depicting the percentage of accessible chromatin fibers were obtained from UCSC’s hs1 trackhub (<u>all.percent.accessible.bw</u>). Genomic coordinates of PHRs present in the five acrocentric chromosomes were obtained from Guarracino *et al.* ^39^

T2T primate genome assemblies were also obtained from the NCBI Genome database with corresponding metadata (centromeric, pericentromeric satellites, long-read methylation levels, and RepeatMasker^84^ annotations) downloaded from the <u>GenomeArk AWS S3 bucket</u>. Cactus 8-way primary progressive alignment between T2T-CHM13v2.0, GRCh38.p14, and the six telomere-to-telomere primate assemblies was obtained from the <u>UCSC Computational</u> <u>Genomics Lab & Platform repository</u> in HAL118 MAF format (<u>8-t2t-apes-2023v2.hal</u>). Whole-genome PhastCons ^80^ conservation scores (<u>approach2.scores_v0.3.bw</u>) and lineage-specific SDs for T2T-CHM13v2.0 and the six primate genomes were obtained from analysis performed by Yoo *et al*. ^26^

All statistically significant potential CTCF-binding sites across all analyzed genomes are publicly available and can be found in **Supplementary Table 1**.

## Acknowledgements

Research reported in this publication was supported by the National Institute of General Medical Sciences of the National Institutes of Health under award number R35GM155468. We would also like to thank Ms Nikol Chantzi for helpful discussions.

