## Supplementary Table 2 for "Comprehensive landscape of potential CTCF-binding sites across the complete telomere-to-telomere human genome"

**Supplementary Table 2: Information about non-human T2T primate assemblies used in this study.**

| Species name | Common name | Version | GenBank accession number |
|---|---|---|---|
| *Gorilla gorilla gorilla* | Western lowland gorilla | v2.0 | GCA_029281585.3 |
| *Pan paniscus* | Pygmy chimpanzee | v2.0 | GCA_029289425.3 |
| *Pan troglodytes* | Chimpanzee | v2.0 | GCA_028858775.3 |
| *Pongo abelii* | Sumatran orangutan | v2.0 | GCA_028885655.3 |
| *Pongo pygmaeus* | Bornean orangutan | v2.0 | GCA_028885625.3 |
| *Symphalangus syndactylus* | Siamang gibbon | v2.1 | GCA_028878055.3 |
