## Supplementary Figure 1 for "Comprehensive landscape of potential CTCF-binding sites across the complete telomere-to-telomere human genome"

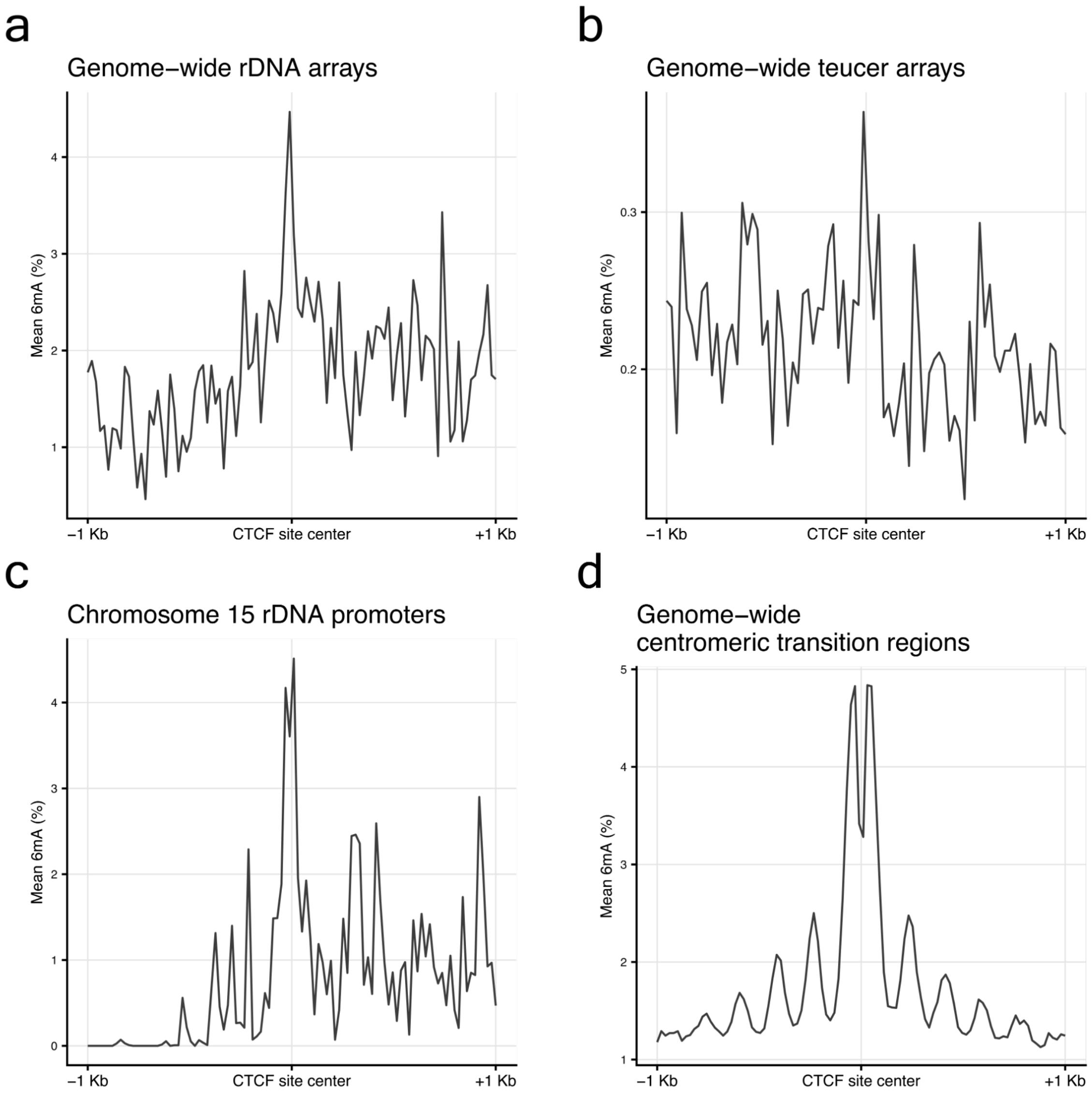


**Supplementary Figure 1: CTCF-directed DiMeLo-seq 6mA metaprofile plots across CTCF-containing genomic regions.** Mean per-bin 6mA from anti-CTCF pA-Hia5 DiMeLo-seq aggregated across CTCF sites and plotted alongside 1 Kb flanks from the motif center. **a.** Genome-wide rDNA regions (5 full-length rDNA arrays; 1,190 potential CTCF sites; 20 bp bins). **b.** Genome-wide teucer arrays (158 regions; 1,179 potential CTCF sites; 25 bp bins). **c.** rDNA promoter regions of chromosome 15 (50 promoters; 100 potential CTCF sites; 20 bp bins). **d.** Genome-wide centromeric transition regions (856 regions; 10,446 potential CTCF sites; 20 bp bins).
