## Supplementary Figure 2 for "Comprehensive landscape of potential CTCF-binding sites across the complete telomere-to-telomere human genome"

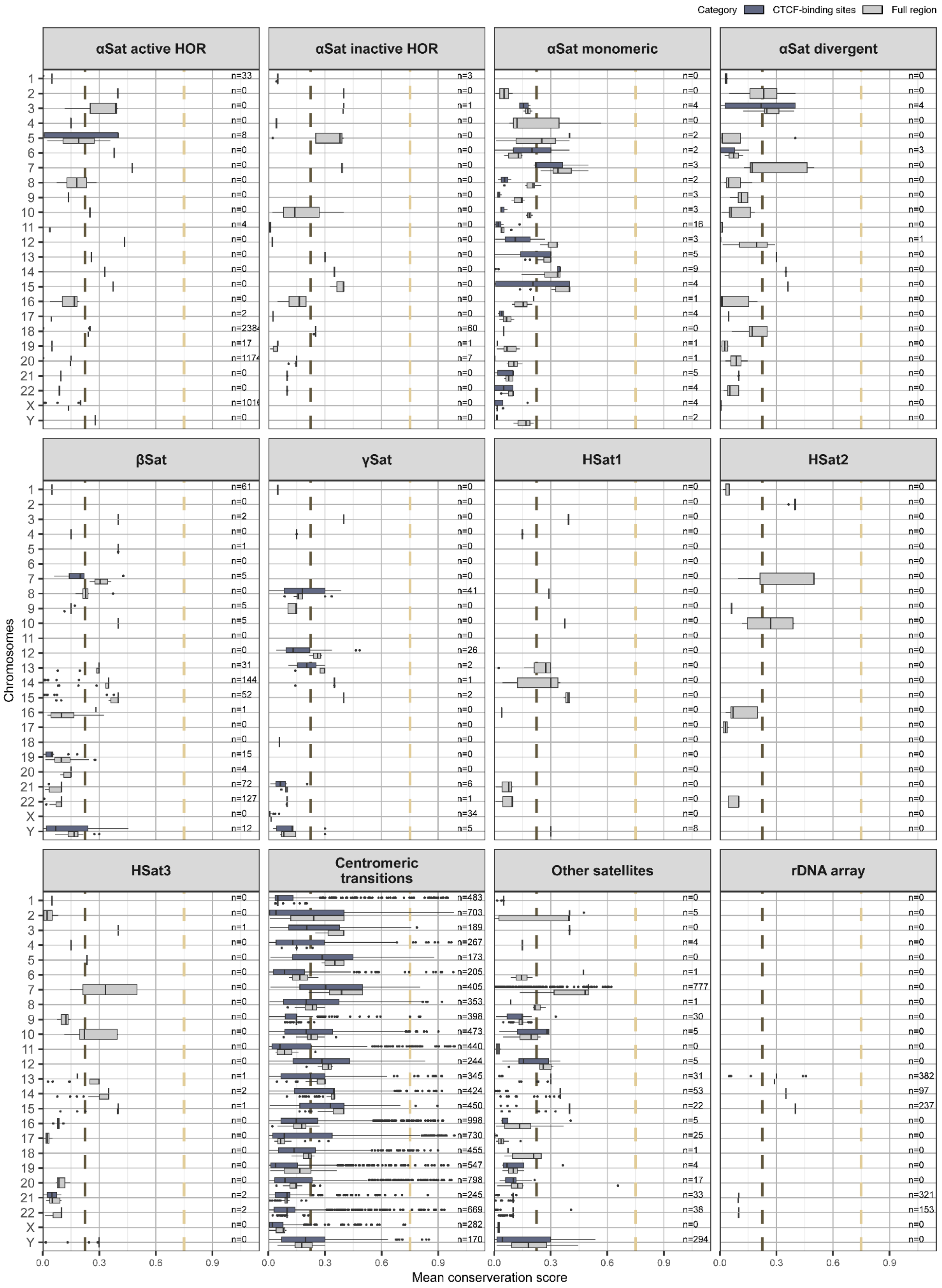


**Supplementary Figure 2: Distribution of CTCF site conservation scores across different CenSat classes.** The grey boxplots represent the conservation of the broader regions, whereas the blue ones represent the conservation of CTCF sites. The brown dashed line represents the mean genome-wide conservation score (score = 0.225), and the gold line represents the threshold used to determine highly conserved CTCF sites (score = 0.75). For each chromosome, n represents the number of CTCF-binding sites found in the chromosome’s satellites.
